# Is there a neural common factor for visual illusions?

**DOI:** 10.1101/2023.12.27.573437

**Authors:** Maya A. Jastrzębowska, Ayberk Ozkirli, Aline F. Cretenoud, Bogdan Draganski, Michael H. Herzog

## Abstract

It is tempting to map interindividual variability in human perception to variability in brain structure or neural activity. Indeed, it has been shown that susceptibility to size illusions correlates with the size of primary visual cortex V1. Yet contrary to common belief, illusions correlate only weakly at the perceptual level, raising the question of how they can correlate with a localized neural measure. In addition, mounting evidence suggests that there is substantial interindividual variability not only in neural function and anatomy but also in the mapping between the two, which further challenges the findings of a neural common factor for illusions. To better understand these questions, here, we re-evaluated previous studies by correlating illusion strengths in a battery of 13 illusions with the size of visual areas and population receptive field sizes. We did not find significant correlations either at the perceptual level or between illusion susceptibility and visual functional neuroanatomy.

## Introduction

Variability is a key element of evolution. Humans are highly idiosyncratic in their appearance, personality traits, cognitive and perceptual abilities, brain anatomy, and physiology. For example, V1 size can vary more than threefold between individuals^1,2^. It is tempting to relate this large variability in neural measures to variability in perception, based on the – often implicit – assumption that perceptual and cognitive processes map onto dedicated brain areas (functional localization). Within this framework, Schwarzkopf and colleagues showed that susceptibility to the Ebbinghaus and Ponzo “hallway” illusions correlates negatively with the surface area of the primary visual cortex V1^3,4^. Similarly, Moutsiana and colleagues reported a significant correlation between size perception in the Delboeuf illusion and population receptive field (pRF) sizes in V1^5^. If localized neural measures, like V1 surface area or pRF size, predict illusion susceptibility, a common factor for illusions should be present at the behavioral level, e.g., higher susceptibility to the Ebbinghaus illusion should go hand in hand with higher susceptibility to the Ponzo illusion. Yet this has not been found to be the case across a multitude of studies. On the contrary, recent studies have shown that visual illusion magnitudes correlate only weakly with each other, even though test-retest reliability is usually good^6–12^. Notably, also in one of the aforementioned studies^3^, V1 surface area was found to correlate with Ebbinghaus and Ponzo illusion magnitudes but the magnitudes of the two illusions did not significantly correlate with each other (*r* = 0.24, *p* = 0.208). Moreover, the correlations between illusion magnitudes and cortical properties were limited to V1 and not found in higher visual areas, i.e., V2, V3 or V4^3–5^, even though surface areas of adjacent areas (e.g., V1 and V2) are known to be highly correlated within individuals^13^.

This multitude of findings is not easy to reconcile. Given the importance of these questions for the general relationship of behavioral and brain measures, our aim here was to replicate previous results. We tested a battery of 13 visual illusions, including the three illusions for which illusion magnitudes have previously been reported to correlate with V1 surface area or pRF size, namely, the Ponzo “hallway”, Ebbinghaus and Delboeuf illusions^3–5^. Additionally, we tested a number of other size illusions, as well as several illusions of perceived orientation, uniform texture, and contrast.

## Results

### Illusion magnitudes

We tested 30 normally-sighted participants (age range: 18 to 35; 14 females) on 13 visual illusions: bisection (BS), contrast (CS), Delboeuf (DB), two variants of the Ebbinghaus (EB1 and EB2), extinction (EX), honeycomb (HC), Müller-Lyer (ML), Poggendorff (PD), two variants of the Ponzo (PZ and PZh), tilt (TT), and Zöllner (ZN) illusions (**Figure 1**). Please see the Methods section for details on each illusion.

**Figure 1.**
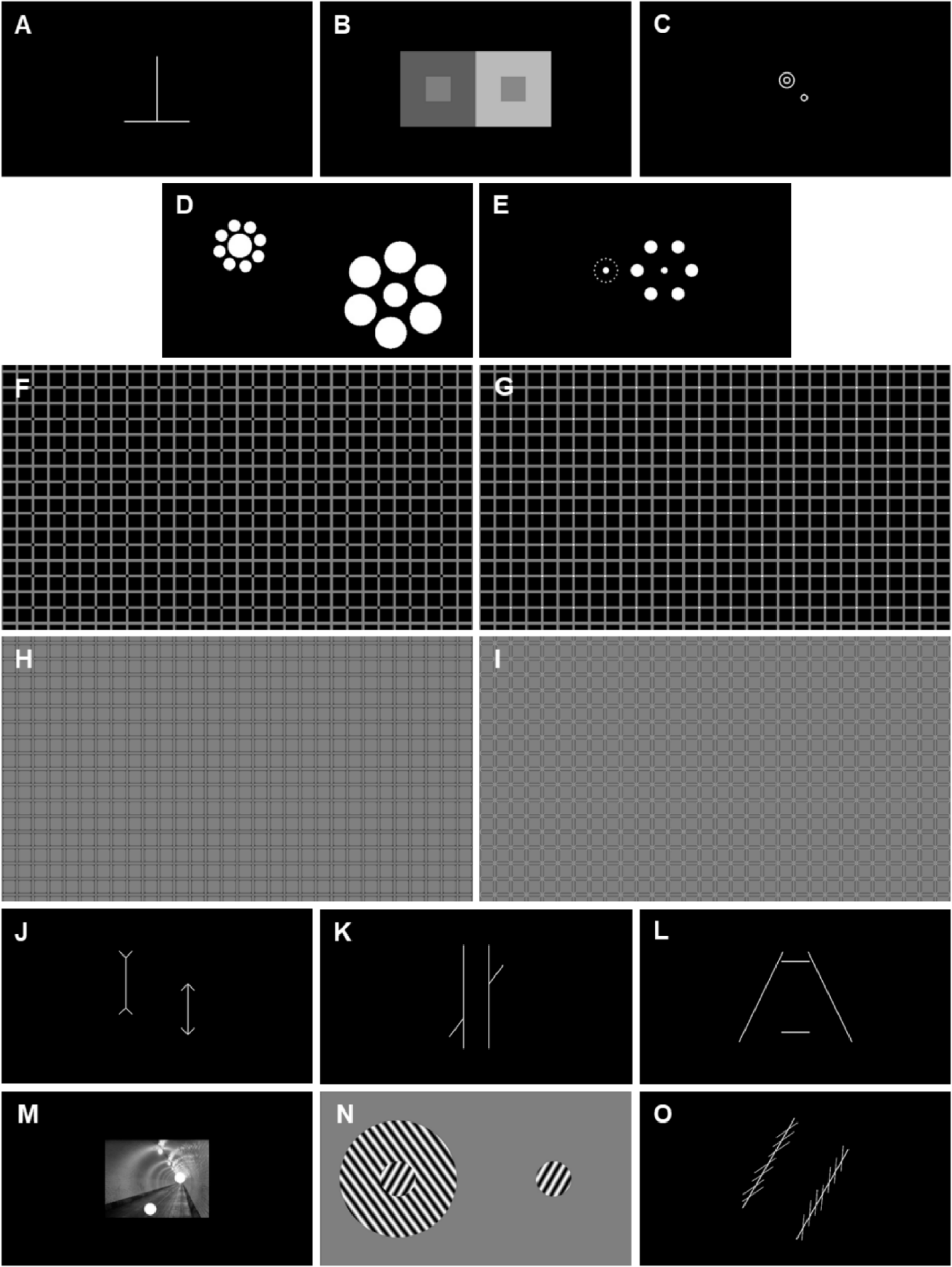
The 13 illusions tested: (A) bisection (BS; participants were instructed to adjust the length of the horizontal segment to match the vertical segment or vice versa), (B) contrast (CS; adjust the shade of gray of the left inner square to match the right inner square or v.v.), (C) Delboeuf (DB; as in previously referenced work^5^; adjust the size of the inner top-left circle to match the bottom-right circle or v.v.), (D) & (E) two variants of the Ebbinghaus (EB1 & EB2, respectively; EB2 as in previously referenced work^4^; adjust diameter of the left central disk to match the right central disk or v.v.), (F) & (G) extinction illusion with black and white dots, respectively (EX; while fixating the center of the screen, adjust the size of the centered red ellipse – not shown here – so that all dots perceived in the periphery of the uniform texture lie within it), (H) & (I) honeycomb illusion with black and white barbs, respectively (HC; while fixating the center of the screen, adjust the size of the centered red ellipse – not shown here – so that all barbs perceived in the periphery of the uniform texture lie within it), (J) Müller-Lyer (ML; adjust length of the left shaft to match the right shaft or v.v.), (K) Poggendorff (PD; adjust the vertical position of the left/right interrupted diagonal so that it lies on a continuum with the right/left interrupted part, respectively), (L) Ponzo (PZ; adjust the length of the upper/lower horizontal segment to match the lower/upper horizontal segment, respectively), (M) Ponzo “hallway” or corridor (PZh; as in previously referenced work^3^; adjust the size of the lower-left disk so that it matches the upper-right disk or v.v.), (N) tilt (TT; adjust the orientation of the right disk to match the left inside disk or v.v.), and (O) Zöllner (ZN; adjust the orientation of one main stream so that it appears parallel to the other main stream) illusions. Illusions (F) to (I) were presented such that they covered a large proportion of the visual field^14^. All illusions except the extinction and honeycomb illusions [(F) to (I)] were tested with two configurations, e.g., in the Ponzo illusion, either the upper horizontal line was adjusted to match the length of the lower horizontal line or v.v.

To test illusion magnitudes, we used an adjustment procedure with the computer mouse. Each illusion was tested with two reference-dependent conditions. For example, in the BS illusion, participants were asked to adjust the length of the vertical segment to match the length of the horizontal segment or vice versa. Participants right clicked the mouse to validate a trial. Each condition was tested twice, with all illusions presented in random order. Unless otherwise specified, illusions were tested under free viewing conditions, i.e., without a fixation point. Illusions were tested in a behavioral testing room, outside of the MRI scanner.

As a measure of test-retest reliability, intraclass correlations were computed between the two adjustments of each condition (see Methods section for details). All ICC coefficients were strong and significant (**Table S1**), except for the contrast illusion (CS left and CS right), which showed small effect sizes.

We calculated illusion magnitudes as the difference compared to the reference, except for the HC and EX illusions, where the area of the adjusted ellipse was considered as a measure of the illusion magnitude. **Figure S1** and **Table S2** show the illusion magnitudes for both conditions of each illusion. All illusions, except the Extinction and Honeycomb illusions, were over-adjusted in one reference-dependent condition and under-adjusted in the other. For example, the horizontal segment of the bisection illusion was over-adjusted when compared to the vertical segment, while the vertical segment was under-adjusted when compared to the horizontal segment.

We combined the two magnitudes of each illusion, i.e., the illusion magnitude of the condition that is usually under-adjusted was added (in absolute values) to the illusion magnitude of the condition that is usually over-adjusted. We then calculated between-illusion correlations (see **Table 1**). In general, most correlations were weak, except for the correlations between the magnitudes of two variants of the same illusion, e.g., between EB1 and EB2, or between PZ and PZh. We also observed a strong correlation between the EX and HC illusion magnitudes, which suggests that a similar mechanism underlies the two illusions, as previously reported^15^.

**Table 1.** Between-illusion correlation coefficients (Pearson’s *r*). Upper triangle: the color scale from blue to red reflects effect sizes from *r* = −1 to *r* = 1. Lower triangle: bold and bold italic font indicate significant results without (*α* = 0.05) and with (*α* = 0.05/78) Bonferroni correction, respectively. Illusion abbreviations: BS – bisection, CS – contrast, DB – Delboeuf, EB1 and EB2 – two variants of the Ebbinghaus, EX – extinction, HC – honeycomb, ML – Müller-Lyer, PD – Poggendorff, PZ and PZh – two variants of the Ponzo, TT – tilt, ZN – Zöllner.

|  | BS | CS | DB | EB1 | EB2 | EX | HC | ML | PD | PZ | PZh | TT | ZN |
| --- | --- | --- | --- | --- | --- | --- | --- | --- | --- | --- | --- | --- | --- |
| BS |  |  |  |  |  |  |  |  |  |  |  |  |  |
| CS | 0.114 |  |  |  |  |  |  |  |  |  |  |  |  |
| DB | -0.280 | 0.066 |  |  |  |  |  |  |  |  |  |  |  |
| EB1 | 0.162 | -0.196 | 0.100 |  |  |  |  |  |  |  |  |  |  |
| EB2 | **0.426** | -0.067 | 0.146 | **0.589** |  |  |  |  |  |  |  |  |  |
| EX | 0.098 | -0.313 | 0.085 | 0.204 | -0.013 |  |  |  |  |  |  |  |  |
| HC | 0.059 | -0.359 | 0.071 | 0.240 | -0.001 | **0.967** |  |  |  |  |  |  |  |
| ML | 0.260 | -0.160 | **-0.428** | 0.048 | 0.029 | -0.126 | -0.015 |  |  |  |  |  |  |
| PD | -0.006 | 0.090 | 0.132 | -0.021 | -0.046 | **0.493** | **0.521** | -0.084 |  |  |  |  |  |
| PZ | -0.156 | -0.083 | 0.142 | 0.221 | 0.053 | 0.268 | 0.348 | 0.114 | 0.219 |  |  |  |  |
| PZh | -0.069 | 0.048 | -0.060 | 0.207 | 0.077 | 0.165 | 0.222 | -0.114 | -0.079 | **0.571** |  |  |  |
| TT | -0.124 | 0.248 | 0.107 | 0.235 | 0.240 | -0.254 | -0.187 | 0.029 | -0.222 | 0.131 | 0.239 |  |  |
| ZN | -0.133 | 0.156 | -0.001 | -0.048 | 0.070 | 0.079 | 0.093 | 0.063 | 0.073 | **0.454** | 0.302 | 0.236 |  |

### Cortical idiosyncrasies and illusion magnitude

All participants completed an MRI session immediately following the behavioral session. The MRI session included six runs of pRF mapping and a high-resolution anatomical scan (see Methods). We used a wedge-and-ring stimulus for pRF mapping, which has been shown to result in a higher model fit and to necessitate shorter acquisition times in comparison with other stimulus configurations^16^. We manually delineated visual areas V1, V2, V3, and V4 on the resulting polar angle maps. We then extracted measures of cortical idiosyncrasy (surface area and pRF size) for each of the visual areas and correlated these measures with the illusion magnitudes obtained for each participant.

#### Visual surface area

Schwarzkopf and colleagues reported negative correlations between retinotopically-defined surface area of the primary visual cortex V1 and Ebbinghaus and Ponzo “hallway” illusion magnitudes^3,4^ and between V1 pRF size and Delboeuf illusion magnitude^5^. To replicate their findings, we first calculated the surface area of visual areas V1, V2, V3, and V4 within eccentricities bounded by the foveal representation and an eccentricity of 8 deg. We then correlated the surface areas with illusion magnitudes from the 13 tested illusions. Here, we group the illusions as follows: (1) illusions previously studied by Schwarzkopf and colleagues (DB, EB2, PZh), (2) other size illusions (BS, EB1, ML, PZ), and (3) uniform texture, perceived orientation, and contrast illusions (CS, EX, HC, PD, TT, ZN). The illusion magnitude-surface area correlations are plotted in separate figures for each of these three groups.

Illusion magnitude-surface area correlations for illusions previously studied by Schwarzkopf and colleagues are shown in **Figure 2**. Bayes Factors in favor of the alternative hypothesis (BF_10_) for each correlation are reported in **Table 2**, along with Pearson’s *r*. We interpreted the strength of evidence in favor of the null or alternative hypothesis based on the following BF_10_ cutoffs^17,18^: substantial evidence for null hypothesis: BF_10_ ∈ [0.1,0.333); anecdotal evidence for null hypothesis: BF_10_ ∈ [0.333,1); anecdotal evidence for alternative hypothesis: BF_10_ ∈ [1,3); substantial evidence for alternative hypothesis: BF_10_ ∈ [3,10); extreme evidence for alternative hypothesis (BF_10_ > 100). In general, our results failed to replicate those reported by Schwarzkopf and colleagues. We did not observe a significant negative correlation between the DB illusion magnitude and V1 surface area, with the Bayes Factor in fact indicating substantial evidence for the null hypothesis (BF_10_ = 0.313). We observed medium effect sizes for the negative correlations in the other ROIs, with a significant correlation in the case of V2 (*p* < 0.05, uncorrected), but the Bayes Factors did not indicate substantial evidence in favor of the alternative hypothesis in any of the ROIs (V2: *r* = −0.373, *p* = 0.043, B_10_ = 1.616; V3: *r* = −0.336, *p* = 0.074, B_10_ = 1.052; V4: *r* = −0.320, *p* = 0.090, B_10_ = 0.905). For EB2, we did not observe any negative correlations between illusion magnitude and visual surface area. In fact, we found (weak) *positive* correlations in visual areas V2 to V4. We also did not observe any significant correlations for the PZh illusion. In fact, the Bayes Factors across all ROIs indicated substantial evidence in favor of the null hypothesis (V1: *r* = 0.005, *p* = 0.979, B_10_ = 0.235; V2: *r* = 0.112, *p* = 0.569, B_10_ = 0.274; V3: *r* = −0.148, *p* = 0.461, B_10_ = 0.309; V4: *r* = 0.128, *p* = 0.526, B_10_ = 0.289).

**Figure 2.**
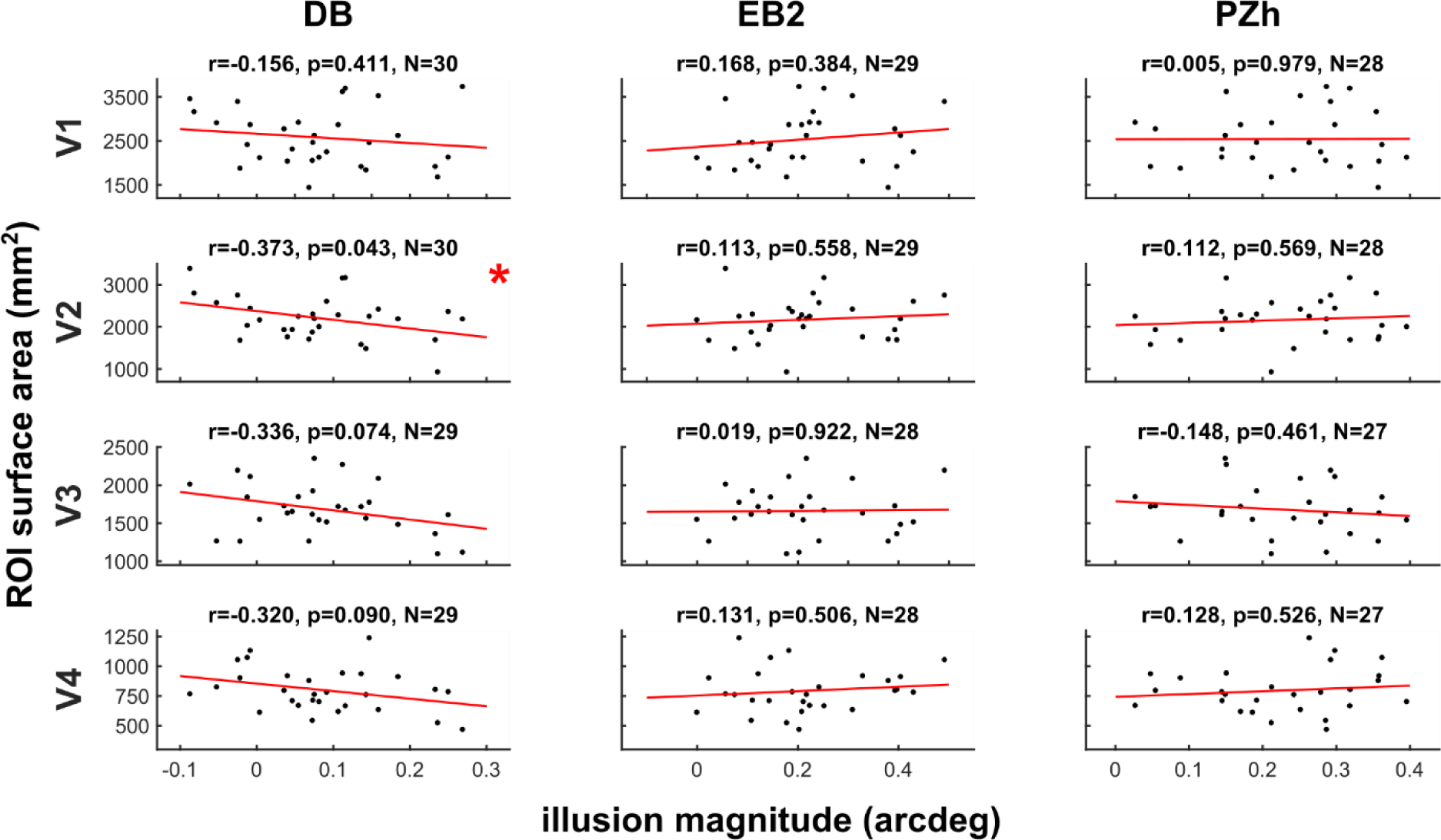
Correlations between ROI surface area and visual illusion magnitudes for illusions previously studied by Schwarzkopf and colleagues. DB – Delboeuf (as in previously referenced work^5^), EB2 – Ebbinghaus variant 2 (as in previously referenced work^4^), PZh – Ponzo “hallway” or corridor (as in previously referenced work^3^). An asterisk marks the one significant correlation (*p* < 0.05, uncorrected). Outliers (absolute modified *z*-score > 3.5) removed.

**Table 2.** Correlations between ROI surface areas (mm^2^) and illusion magnitudes. Significant correlations (*p* < 0.05, uncorrected) are marked in bold with an asterisk. Only one of these correlations constitutes at least substantial evidence in favor of the alternative model. In fact, this correlation – between V3 surface area and BS illusion magnitude – constitutes extreme evidence in favor of the alternative model (BF_10_ > 100) and is marked in bold with three asterisks. Instances of substantial evidence in favor of the null hypothesis (BF_10_ ∈ [0.1,0.333)) are marked in red and bold with a dagger (†). Outliers (absolute modified *z*-score > 3.5) removed.

| Visual area | Illusion | N | Pearson's $r$ | $BF_{10}$ |
| --- | --- | --- | --- | --- |
| V1 | BS | 30 | 0.317 | 0.912 |
|  | CS | 30 | 0.189 | 0.366 |
|  | DB | 30 | -0.156 | **0.313†** |
|  | EB1 | 30 | 0.170 | 0.334 |
|  | EB2 | 28 | 0.032 | **0.238†** |
|  | EX | 29 | -0.172 | 0.337 |
|  | HC | 30 | -0.074 | **0.244†** |
|  | ML | 29 | 0.168 | **0.331†** |
|  | PD | 29 | -0.251 | 0.526 |
|  | PZ | 30 | -0.101 | **0.259†** |
|  | PZh | 28 | 0.005 | **0.235†** |
|  | TT | 30 | 0.225 | 0.449 |
|  | ZN | 30 | -0.132 | **0.286†** |
| V2 | BS | 30 | 0.360 | 1.397 |
|  | CS | 30 | 0.122 | **0.277†** |
|  | DB | 30 | **-0.373\*** | 1.616 |
|  | EB1 | 30 | 0.211 | 0.413 |
|  | EB2 | 28 | 0.051 | **0.242†** |
|  | EX | 29 | -0.164 | **0.326†** |
|  | HC | 30 | 0.074 | **0.244†** |
|  | ML | 29 | 0.113 | **0.272†** |
|  | PD | 29 | -0.343 | 1.127 |
|  | PZ | 30 | -0.138 | **0.292†** |
|  | PZh | 28 | 0.112 | **0.274†** |
|  | TT | 30 | 0.209 | 0.407 |
|  | ZN | 30 | -0.248 | 0.523 |
| V3 | BS | 29 | **0.628\*** | **130.580\*\*\*** |
|  | CS | 29 | 0.136 | **0.292†** |
|  | DB | 29 | -0.336 | 1.052 |
|  | EB1 | 29 | 0.185 | 0.358 |
|  | EB2 | 27 | 0.016 | **0.240†** |
|  | EX | 28 | -0.018 | **0.236†** |
|  | HC | 29 | 0.105 | **0.265†** |
|  | ML | 28 | 0.019 | **0.236†** |
|  | PD | 28 | **-0.386\*** | 1.656 |
|  | PZ | 29 | -0.200 | 0.386 |
|  | PZh | 27 | -0.148 | **0.309†** |
|  | TT | 29 | -0.104 | **0.265†** |
|  | ZN | 29 | **-0.419\*** | 2.633 |
| V4 | BS | 29 | 0.272 | 0.611 |
|  | CS | 29 | 0.010 | **0.231†** |
|  | DB | 29 | -0.320 | 0.905 |
|  | EB1 | 29 | 0.158 | **0.318†** |
|  | EB2 | 27 | 0.048 | **0.245†** |
|  | EX | 28 | -0.037 | **0.239†** |
|  | HC | 29 | -0.136 | **0.292†** |
|  | ML | 28 | 0.131 | **0.290†** |
|  | PD | 28 | -0.201 | 0.387 |
|  | PZ | 29 | -0.198 | 0.383 |
|  | PZh | 27 | 0.128 | **0.289†** |
|  | TT | 29 | -0.045 | **0.237†** |
|  | ZN | 29 | -0.048 | **0.237†** |

**Figure S1** shows the illusion magnitude-surface area correlations for the other size illusions (BS, EB1, ML, PZ). Here, we did not observe any significant negative correlations. We observed weak negative correlations for the ML and PZ illusions across ROIs, but none of them was significant. Contrary to the observation of negative correlations between V1 surface areas and size illusion magnitudes by Schwarzkopf and colleagues, we observed *positive* correlations for the BS illusion across visual areas, though the correlation was significant only in V3 (*r* = 0.628, *p* = 2.62e-4), with “extreme” evidence (BF > 100)^17,18^ in favor of the alternative hypothesis (BF_10_ = 130.580). Schwarzkopf and colleagues suggested that the mediating factor between illusion magnitude and cortical size is the spatial spread of neuronal connections between the target and inducer. Thus, the finding of a positive correlation is surprising since it suggests that larger visual cortical surface areas (to a limited extent) give rise to stronger BS illusion susceptibility. Likewise, we observed weak positive correlations for EB1 across all ROIs, though none of these correlations was significant.

Illusion magnitude-surface area correlations for all other illusions (uniform texture, perceived orientation, contrast) are shown in **Figure S2**. We observed two significant correlations between illusion magnitude and visual surface area, one for the PD illusion in V3 (*r* = −0.386, *p* = 0.043, BF_10_ = 1.656) and another for ZN in V3 as well (*r* = −0.419, *p* = 0.024, BF_10_ = 2.633), but neither of these constituted substantial evidence in favor of the alternative hypothesis.

#### PRF size and illusion magnitude

Previous studies have investigated the role of macroscopic visual surface area in determining interindividual variability in perception under the assumption that it is a proxy for local variations in cortical magnification or, inversely, pRF size. The initial seminal studies linking illusion susceptibility with early visual surface areas^3,4^ relied on phase-encoded methods rather than model-based^19^ retinotopic mapping, which – though it allowed for a manual delineation of early visual areas based on reversals in the polar angle map – did not allow for the estimation of pRF size. Here, we used pRF modeling to directly investigate the role of pRF size in determining illusory size perception. Moreover, we sought to replicate the findings from the other previously mentioned study by Moutsiana and colleagues^5^, which used model-based pRF mapping and reported a negative correlation between pRF size and Delboeuf illusion magnitude. To this end, we extracted pRF sizes from eccentricities relevant to each given illusion and correlated them with illusion magnitudes.

This analysis could only be done for illusions in which the targets are at a fixed distance from one another. Moreover, the distance between the targets could be reasonably determined only for circular targets. Thus, we focused on the DB and EB2 illusions. In both cases, we assumed that participants used one of two strategies: (1) focusing their gaze at the midpoint location between the two targets, or (2) focusing their gaze directly on one or the other of the targets. We extracted pRF sizes from illusion-relevant eccentricities in accordance with both strategies (see Methods for details). From this point forward, we refer to the eccentricities used in strategy (1) as the “larger eccentricity” and strategy (2) as the “smaller eccentricity”.

Correlations between illusion magnitudes and pRF sizes at illusion-relevant eccentricities are shown in Figure 3 for the Delboeuf illusion and in Figure 4 for the Ebbinghaus 2 illusion. The Bayes Factor in favor of the alternative hypothesis (BF_10_) for each correlation is reported in **Table 3**, along with Pearson’s *r*. In general, we failed to observe an association between pRF size and illusion magnitude. For the Delboeuf illusion, we observed two medium effect size *positive* correlations in V3, one in the case of the smaller eccentricity (1.96 deg: *r* = 0.341, *p* = 0.066, B_10_ = 1.145) and the other in the case of the larger eccentricity (3.92 deg: *r* = 0.360, *p* = 0.051, B_10_ = 1.404). Neither of the two was significant and constituted only anecdotal evidence in favor of the alternative hypothesis. Similarly, for the Ebbinghaus 2 illusion, we observed two correlations of medium effect size in V3 (*r* = 0.319, *p* = 0.099, BF_10_ = 0.861) and V4 (*r* = 0.444, *p* = 0.016, BF_10_ = 3.697), both for the smaller eccentricity (4.65 deg). Only the latter correlation was significant, with the Bayes factor indicating substantial evidence in favor of the alternative hypothesis.

**Figure 3.**
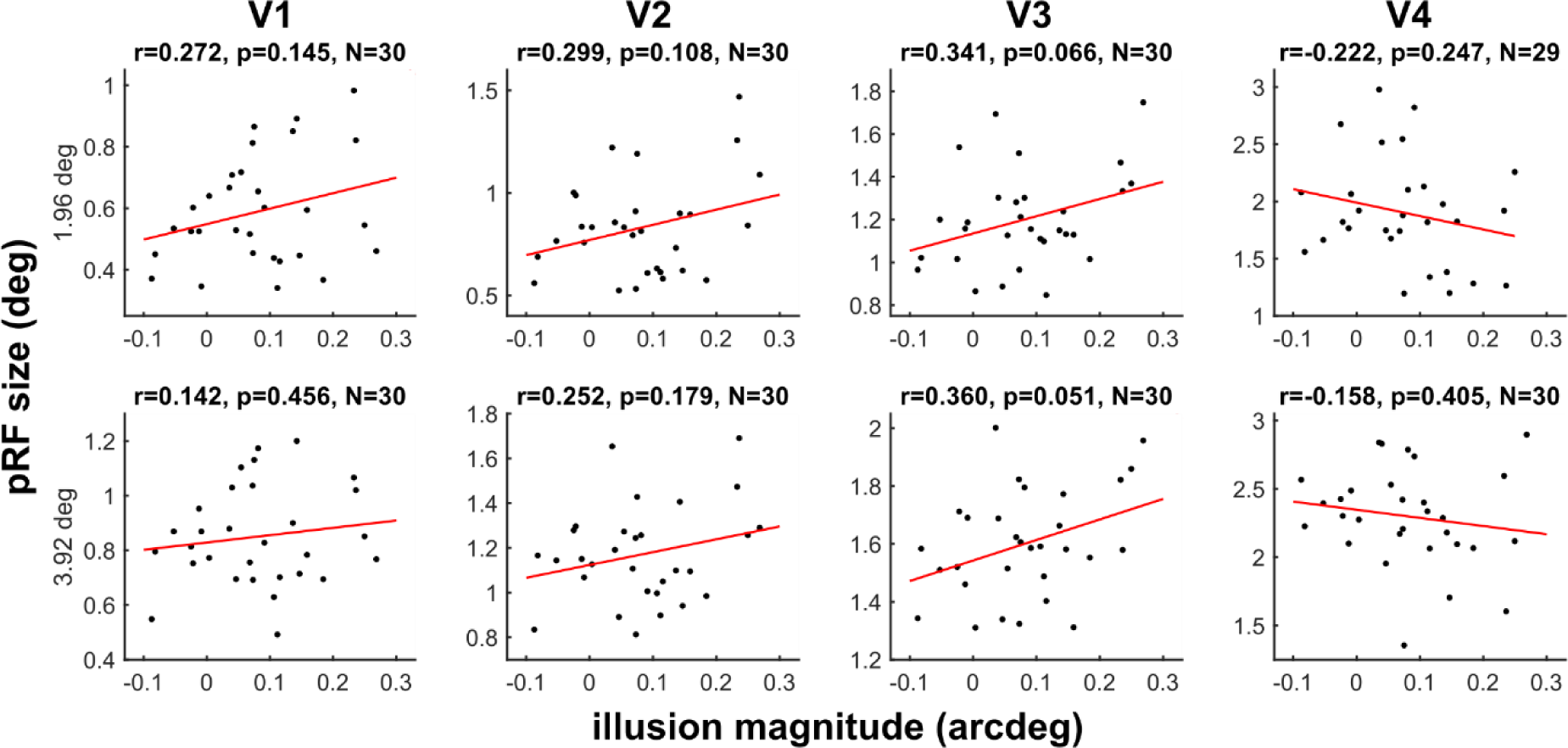
Correlations between pRF size at illusion-relevant eccentricities and visual illusion magnitudes for the Delboeuf illusion. None of the correlations are significant (*p* < 0.05, uncorrected).

**Figure 4.**
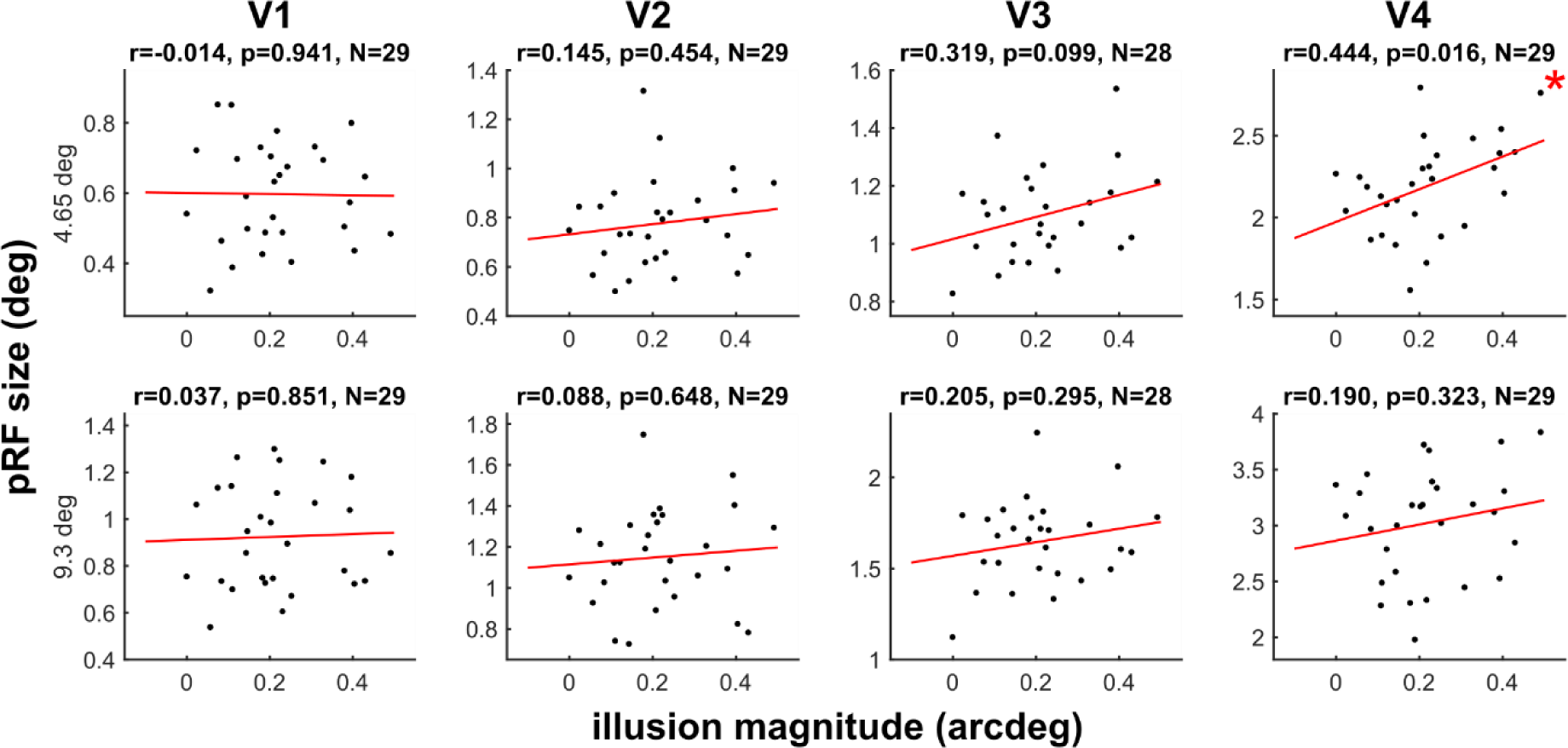
Correlations between pRF size at illusion-relevant eccentricities and visual illusion magnitudes for the Ebbinghaus 2 illusion. An asterisk marks the one significant correlation (*p* < 0.05, uncorrected for multiple comparisons).

**Table 3.** Correlations between pRF size (deg) at relevant eccentricities and illusion magnitudes with outliers removed. The one significant correlation (*p*<0.05, uncorrected) is marked in bold with an asterisk. Only this single correlation constitutes substantial evidence in favor of the alternative model (BF_10_ ∈ [3,10)). It is marked in bold with an asterisk. Numerous correlations constitute substantial evidence in favor of the null hypothesis (BF_10_ ∈ [0.1,0.333)) – these are marked in red and bold with a dagger (†).

| Visual area | Illusion | Eccentricity (deg) | n | Pearson's $r$ | $BF_{10}$ |
| --- | --- | --- | --- | --- | --- |
| V1 | DB | 1.96 | 30 | 0.272 | 0.625 |
|  |  | 3.92 | 30 | 0.142 | **0.296†** |
|  | EB2 | 4.65 | 29 | -0.014 | **0.231†** |
|  |  | 9.3 | 29 | 0.037 | **0.235†** |
| V2 | DB | 1.96 | 30 | 0.299 | 0.779 |
|  |  | 3.92 | 30 | 0.252 | 0.538 |
|  | EB2 | 4.65 | 29 | 0.145 | **0.302†** |
|  |  | 9.3 | 29 | 0.088 | **0.255†** |
| V3 | DB | 1.96 | 30 | 0.341 | 1.145 |
|  |  | 3.92 | 30 | 0.360 | 1.404 |
|  | EB2 | 4.65 | 28 | 0.319 | 0.861 |
|  |  | 9.3 | 28 | 0.205 | 0.395 |
| V4 | DB | 1.96 | 29 | -0.222 | 0.438 |
|  |  | 3.92 | 30 | -0.158 | **0.316†** |
|  | EB2 | 4.65 | 29 | **0.444\*** | **3.697\*** |
|  |  | 9.3 | 29 | 0.190 | 0.368 |

#### Slope and intercept of pRF size as a function of eccentricity

The above pRF analysis was limited to just two illusions due to the difficulty of identifying “relevant” eccentricities for most illusions. To circumvent this issue and characterize the relationship between pRF size and illusion magnitude for all tested illusions, we estimated the slope and intercept of pRF size as a function of eccentricity. An association between the slope and illusion magnitude would indicate that interindividual susceptibility to the given illusion is predicated on the rate at which pRF size increases with eccentricity. Meanwhile, an association between the intercept and illusion magnitude would imply that the parafoveal pRF size is the determining factor for illusion susceptibility.

For each participant, each ROI, and the two hemispheres separately, we fit a linear regression model to the pRF size as a function of eccentricity, weighted by the pRF model’s goodness-of-fit (R^2^) at each given eccentricity. We calculated the slope and intercept of the regression line and averaged the slopes and intercepts from the two hemispheres (for further details see the Methods section).

For each illusion and each ROI, we conducted Bayesian linear regression analyses, with illusion magnitudes as the dependent variable and slope and/or intercept as the covariate(s). We tested four models: (1) a model that included the slope, (2) a model that included the intercept, (3) a model that included both slope and intercept, (4) a null model that included neither slope nor intercept. Bayes Factors (BF_10_) in favor of each of the three alternative Bayesian linear regression models – slope, intercept, and slope + intercept – are reported in **Table S3**. We did not find substantial evidence for any of the tested models with respect to the null model for any of the tested illusions. For most of the other slope and intercept models, we found anecdotal evidence for the null hypothesis. Meanwhile, for the model containing both the slope and intercept terms, we found substantial evidence in favor of the null model for most illusions and ROIs (BF_10_ ∈ [0.1,0.333)).

## Discussion

One of the main aims of cognitive neuroscience is the mapping of perceptual and cognitive functions onto the brain^20^. Cognitive neuroscience studies often make the implicit assumption that functions can be attributed to localized brain areas (as opposed to distributed networks). Here, we considered the example of visual illusions, with the aim of verifying previous reports of a localized neural predictor of size illusion magnitude^3–5^.

Intuitively, one might expect that someone with high susceptibility to the Ebbinghaus illusion has high susceptibility to the Ponzo illusion as well. Taxonomizing illusions as spatial, temporal, and contrast illusions reflects this thinking. Recent research has clearly shown that strong correlations between different illusions are rare^6–12^ (but see the recent work of Makowski and colleagues^21^). In fact, Schwarzkopf and colleagues also reported that the illusion magnitudes in the Ebbinghaus and Ponzo illusions did not correlate significantly^3^. The low correlations cannot be explained by low test-retest reliability, lack of power, or a limited range in the illusion magnitudes^6,9,10^. Factor analyses support the notion of low correlations and suggest that each illusion makes up its own factor^6–8,22^. It must be noted that findings of correlations between illusion susceptibility and neural measures while at the same time inter-illusion correlations are weak could be sound, as correlations are only transitive when they are sufficiently close to 1^23^. However, to claim that individual variation in brain area X *predicts the performance* of visual task Y, where visual task Y is a category of illusions, these findings should generalize at least across illusions of the given category.

To resolve these conceptual misgivings, we conducted a systematic study with a variety of illusions. We could not reproduce previous findings of negative correlations between early visual surface areas and susceptibility to different size illusions (DB, EB2 and PZh), as reported by Schwarzkopf and colleagues. Almost none of the correlations were significant. While we observed *negative* correlations for some selected illusions and brain regions (small-to-medium effect sizes), the rest of the correlations were mostly *positive* (small-to-medium effect sizes). In addition to the surface area analysis, we tested whether illusion susceptibility can be explained by pRF size (as in previously referenced work^5^) for DB and EB2. We failed to replicate previous findings, observing no significant correlations. In a final analysis, we tried to explain illusion susceptibility with the parafoveal pRF size and the rate of change of pRF size from fovea to periphery – using the intercept and slope of the linear relation between pRF size and eccentricity. Once again, we did not find evidence in support of the claim of localized neural measures explaining interindividual differences in illusion magnitudes.

Our study goes well beyond the previously mentioned studies^3–5^. First, we tested a much larger battery of visual illusions – 13 in total. These included the illusions tested in the referenced literature – Delboeuf, Ebbinghaus and the Ponzo hallway illusion – but also four other size illusions – bisection, another variant of Ebbinghaus, Müller-Lyer and Ponzo – and six other illusions of uniform texture, perceived orientation, and contrast. This large batch allowed us to test the replicability and generalizability of previously mentioned results. A second important difference was that the aforementioned studies used phase-encoded methods to functionally-define visual surface areas as a measure of central visual cortical magnification^3,4^. Here, we used model-based pRF mapping^19^, which, in addition to providing more robust eccentricity measures, also estimates pRF size, i.e., the spatial spread of neural activation in response to the visual stimulus. This allowed us to correlate illusion magnitudes not only with V1 surface area but also with pRF size, as in previously referenced work^5^. Thirdly, in our study we used free viewing conditions, which meant that we did not control participants’ subjective experience of the illusions due to individual gaze patterns. As a result, we had to make assumptions about the viewing strategies participants used. To account for any systematic biases that can result from the assumptions of the two strategies, future studies should include a fixation point. Fourthly, we used an adjustment task, whereas the aforementioned studies used psychometric curve fitting and a Multiple Alternative Perceptual Search (MAPS) task^3–5^. However, we have previously shown that individual differences are stable across different measurement methods^24^, and test-retest reliability was high across all tested illusions.

There are increasing calls for large cohort studies or individual-focused precision studies due to reproducibility and statistical power concerns^25–27^. Indeed, statistical simulations show that correlation measures are highly unstable at low sample sizes^28^. Here, we had a sample size of 30 participants, which was the same as in the first of the previously cited work by Schwarzkopf and colleagues^3^ and higher than in the other studies mentioned above (*N* = 26^4^; *N* = 10^5^). While this constitutes an above-average sample size in neuroimaging studies, it is not sufficient to detect small effects. Schwarzkopf and colleagues reported medium correlations between V1 surface area and Ebbinghaus (*r* = −0.38) and Ponzo (*r* = −0.48) illusion magnitudes^3^, which, assuming a significance level of *α* = 0.05, corresponds to achieved power of 0.560 and 0.790, respectively. Furthermore, in the work of Moutsiana and colleagues^5^, the sample size was 10, but the authors treated the four dependent data points from the four visual quadrants of each participant as independent data in their statistical analyses, ending up with an artificially inflated sample size of 40. These sampling issues could partially explain our failure to replicate previous findings.

The quest for neural predictors of cognitive functions, i.e., functional localization, lies at the heart of cognitive neuroscience. While the scope of functional localizationism – whether the location is constrained to a single brain region or distributed in networks – has been actively debated for nearly two centuries^29–33^, another fundamental question is how to define the “function” that we are trying to map. The cognitive ontology that one presumes determines whether and where a selective structure-function mapping is found^20,34^. Traditionally, cognitive ontologies have been constructed through the observance of common factors in personality, cognitive ability, behavior, and perception. The way in which illusions are ontologized is critical in identifying whether robust neural explanations exist; different illusions or illusion categories might have different neural substrates. It is also unclear what neural substrate one should consider in the first place. Schwarzkopf and colleagues mapped illusion magnitude onto V1 size, but it is equally conceivable that another brain metric, like gray matter volume or alpha neural oscillations, might also correlate. Indeed, Schwarzkopf and colleagues have shown evidence for such correlations^35–37^. Moreover, neuroscientists usually assume that brain structure-function mappings are consistent across individuals^38,39^. Yet increasing evidence suggests that in addition to variability in neural function and neuroanatomy, humans vary substantially in the mapping between the two, with different neuronal systems capable of carrying out the same functions^40–42^. Our results show that the current approach to categorizing illusions and searching for a selective function-structure mapping does not always lead to reproducible results.

In conclusion, our results speak against the existence of (localized) neural common factors for visual illusions. Our study goes beyond a failure to replicate, showing that the deeply engrained view about common factors does not seem to hold for visual illusions. The large variability in both neural and perceptual measures, with low correlations at the perceptual level, makes it unlikely that there are simple links between variability on one and the other level. We advocate for the need to shift studies of interindividual variability to more sophisticated neuroimaging data acquisition and analysis methods, e.g., predicting pattern of visual illusion results based on patterns in brain data^38^. Constructing a more realistic ontology of visual cognition is critical to successfully describing the brain structure-function relationship and verifying the empirical validity and nature of functional localization.

## Methods

### Participants

Thirty paid volunteers (14 females), 18 to 35 years of age (mean ± standard deviation: 24.7±4.45) participated in the experiment. All but six were right-handed. All participants had normal uncorrected visual acuity – both binocular and monocular in both eyes – as assessed with the Freiburg Visual Acuity test, i.e., acuity values above 1.0^43^. The experiment included a psychophysics part – carried out in a behavioral testing room – and an fMRI part, which immediately followed the psychophysics part. Participants gave written informed consent and were made aware that they could discontinue the experiment at any time. All experimental procedures complied with the Declaration of Helsinki, except for pre-registration (§35), and were approved by the local ethics committee. All participants were naïve to the purpose of the experiment.

### Stimuli and apparatus

For the psychophysics experiment, visual stimuli were displayed on an ASUS VG278HE LCD monitor (refresh rate: 60 Hz; spatial resolution: 1920×1080) at a viewing distance of 76 cm. The mean luminance of the screen was 75 cd/m^2^. Stimulus programs were implemented in MATLAB (The MathWorks, Inc., Natick, MA) using the Psychtoolbox^44^.

The MRI experiment was conducted at the MRI-scanning facilities of the Department of clinical neurosciences at the Centre Hospitalier Universitaire Vaudois (CHUV) – Lausanne, Switzerland. Participants lay in the MRI scanner and looked at a screen, placed at the end of the 60 cm scanner bore, through a head-coil mounted mirror (viewing distance: 70 cm). A Sony VPL-FH31 projector (size of projected image: 57×32.1 cm, chosen pixel resolution: 1920×1080 pixels, refresh rate: 60 Hz) was used to back-project the stimuli onto the screen, with a mean luminance of 1000 cd/m^2^. As in the psychophysics part, stimulus generation and response collection were done in MATLAB using the Psychtoolbox.

#### Illusion stimuli

Thirteen illusions were tested (Figure 1) using an adjustment procedure: the bisection (BS), contrast (CS), Delboeuf (DB), two variants of the Ebbinghaus (EB1 and EB2), extinction (EX), honeycomb (HC), Müller-Lyer (ML), Poggendorff (PD), two variants of the Ponzo (PZ and PZh), tilt (TT), and Zöllner (ZN) illusions. Each illusion was tested with two reference-dependent conditions, e.g., participants were asked to adjust the length of the vertical segment to match the length of the horizontal segment in the BS illusion, or vice versa. Participants performed the adjustment with a computer mouse and then right clicked to validate the trial. Each condition was tested twice, making up 52 trials in total (13 illusions × 2 reference-dependent conditions × 2 trials), presented in a random order. No feedback was provided and there was no time limit, i.e., the stimuli were shown until a response was given. Unless otherwise specified, a free viewing condition was used, i.e., there was no fixation point. Lines were shown with a 4-pixel width, except in the DB illusion, where the circles were 2 pixels wide.

##### Bisection (BS; also called vertical-horizontal)

Participants had to adjust the length of the vertical segment so that it appeared to be the same length as the horizontal segment or to adjust the length of the horizontal segment to match the length of the vertical one (Figure 1A). The reference segment was 10.4° long and the adjustable segment had a size randomly set in the range of 2-17° at the beginning of each trial. The horizontal segment was displayed 5.2° to the bottom of the middle of the screen and was always touching the vertical segment.

##### Contrast (CS)

We asked participants to adjust the shade of gray of the left inside square so that they perceived it to be the same as the shade of gray of the right inside square, or vice versa (Figure 1B). The sides of the inside squares were 4° in length. The inside squares were displayed in the middle of two outside squares, whose sides were 12° in length. The entire inside-outside square configuration was displayed in the middle of the screen.

##### Delboeuf (DB)

The stimulus was strongly inspired by previously referenced work^5^, where the authors used a Multiple Alternative Perceptual Search (MAPS) procedure, in which participants had to report which of four comparison rings was the more similar in size to a central target. Here, we used an adjustment procedure, in which participants had to adjust the size of the upper-left inside circle to match the size of the lower-right circle or vice versa (Figure 1C). The reference and upper-left outside circles were 0.98° and 2.35° in diameter, respectively. The size of the adjustable circle was randomly set at the beginning of each trial, but it never exceeded 2.35°. The reference and adjustable circles were 3.92° from each other and the whole illusion was centered in the middle of the screen.

##### Ebbinghaus (EB1 and EB2)

In both variants of the Ebbinghaus illusion, participants were instructed to adjust the size of the left central disk (i.e., left target) to match the size of the right central disk (i.e., right target), or vice versa.

We measured the susceptibility to the Ebbinghaus illusion (EB1) as we did previously^6^. The left and right targets were surrounded by eight small (2.0° in diameter) and six large (5.2° in diameter) flankers, respectively (Figure 1D). The reference target was 4° in diameter. The centers of the small and large flankers were 3.36° and 6.08° away compared to the center of the left and right targets, respectively. The left and right targets were centered 12.41° to the left and right and 4° to the top and bottom, respectively, compared to the center of the screen.

The second variant of the Ebbinghaus illusion (EB2) was strongly inspired by previously referenced work^4^. The left and right targets were surrounded by 16 small (0.26° in diameter) and six large (2.07° in diameter) flankers, respectively (Figure 1E). The small and large flankers were located 1.86° and 4.34° away from the center of the left and right targets, respectively. The reference target was 1.03° in diameter. The left and right targets were located at 4.65° eccentricity compared to the middle of the screen and were vertically centered.

In both variants of the Ebbinghaus illusion, the size of the adjustable target was randomly chosen by the computer at the beginning of each trial with the constraint that it could never touch the flankers.

##### Extinction (EX) and Honeycomb (HC)

The stimuli used here were the same as in previously referenced work^14,45^. In the Honeycomb (HC) and Extinction (EX) illusions, participants are unable to see shapes (barbs in the HC illusion; dots in the case of the EX illusion) in the periphery of a uniform texture, while they are fixating the center of the texture.

Participants were asked to fixate a red central cross while adjusting the size of a red ellipse on both *x* and *y* dimensions, so that they could perceive all barbs (HC) and dots (EX) inside the ellipse. The initial size of the red ellipse was randomly chosen at the beginning of each trial, with the screen size as limits. Two contrast polarity conditions, i.e., black or white barbs (HC) or dots (EX), were tested for each illusion, making up four conditions (Figure 1F-I). To reduce the aftereffect following a HC or EX trial, 30 random light and dark gray checkerboards made of squares (0.52° length of each side) with 0.35 and 0.65 of the maximum luminance were presented for 0.5 second each.

##### Müller-Lyer (ML)

Participants had to adjust the length of the left shaft (i.e., the vertical segment with inward-pointing arrows) so that they perceived it as long as the right shaft (i.e., the vertical segment with outward-pointing arrows), or vice versa (Figure 1J). The reference shaft was 8° long and the fins were 1.5° long, oriented at 45° compared to the vertical. At the beginning of each trial, the length of the adjustable shaft was randomly set between 2 and 21°. The left and right shafts were centered 2.12° to the top and bottom and 4.97° to the left and right compared to the middle of the screen, respectively.

##### Poggendorff (PD)

Participants were instructed to adjust the vertical position of the left or right interrupted diagonal so that it appeared to lie on a continuum with the right or left interrupted part, respectively (Figure 1K). The two main vertical streams were 16.6° long and 4° away from each other. Both parts of the interrupted diagonal were 3.8° long and titled by 37° compared to the vertical. When the position of the left part of the interrupted diagonal was adjusted, the right part was touching the right main stream 6.29 degrees away compared to the top of the main streams. When adjusting the position of the right part of the interrupted diagonal, the left part touched the left main stream 5 degrees away compared to the bottom of the main streams. The adjustable element was randomly positioned along the corresponding main stream at the beginning of each trial and the whole illusion was centrally displayed.

##### Ponzo (PZ) and Ponzo “hallway” (PZh; also called corridor)

In the first variant of the Ponzo (PZ) illusion, the task was to adjust the length of the upper or lower horizontal segment to match the length of the lower or upper horizontal segment, respectively (Figure 1L). The reference segment was 4.5° long, while the length of the adjustable segment was randomly set between 0 and 12° at the beginning of each trial. To induce a trapezoid-like perspective in the illusion, two converging lines were shown with 4° separating them at the apex and 18° at the base. The total height of the imaginary trapezoid was 14.4° and the reference and adjustable segments were 11.3° away from each other. The whole illusion was centrally displayed.

The second variant of the Ponzo (PZh) illusion was inspired by previously referenced work^3^. Participants were instructed to adjust the size of the lower-left disk so that it appeared to be the same size as the upper-right disk or vice versa (Figure 1M). A tunnel image (640 × 480 pixels) was displayed in the center of the screen on a black background. The reference disk was 1° in diameter and the size of the adjustable disk was set in the range of 0 to 2° in diameter at the beginning of each trial. The adjustable disk was fixed at its lowest point, as if it was anchored to the image background.

##### Tilt (TT)

Participants had to adjust the orientation of the right disk to match the orientation of the left inside disk or vice versa (Figure 1N). The reference and adjustable disks were 6° in diameter and made of a 0.5 cycles/° full contrast grating texture. The reference disk was titled 33° clockwise compared to the vertical and displayed with a random orientation at the beginning of each trial. The outside left disk was 20° in diameter and titled 36° counterclockwise compared to the vertical (0.5 cycles/° full contrast grating texture). The left and right disks were displayed 6.98° to the left and right compared to the middle of the screen.

##### Zöllner (ZN)

Participants were instructed to adjust the orientation of one main stream so that it appeared parallel to the other main stream (Figure 1O). The main streams were 16.6° long, 9.94° apart from each other and tilted by 30° compared to the vertical. At the beginning of each trial, the adjustable main stream was randomly tilted between 0 and 90° compared to the vertical. Seven segments, 4.15° long, were intersecting each main stream. They were tilted 25° with respect to the main streams and their position relative to the main stream was randomly shifted between ±0.83°.

#### Population receptive field mapping stimulus

The pRF mapping procedure was based on the one suggested by Alvarez and colleagues to yield comparatively high pRF model fit among other commonly used stimulus configurations^16^. The stimulus consisted of a simultaneous rotating wedge and expanding and contracting ring (**Figure S5**). The dimensions of the wedge and ring as well as the timing parameters were very similar to those described by van Dijk and colleagues^46^. Each session ended with a 45-s period of fixation, with a total of 235 volumes acquired. Please see the Supplementary Information for details.

### MRI data acquisition

Participants completed an MRI session directly following the behavioral session. MRI data were acquired using a 3 T whole-body MRI system (Magnetom Prisma, Siemens Medical Systems, Germany), using a 64-channel RF receive head coil and body coil for transmission. A high-resolution anatomical image (T1-weighted MPRAGE, 1.0 mm isotropic resolution) was acquired first and used to place the bounding boxes for the remaining sequences. Participants completed a practice run of the fMRI tasks during this acquisition. Functional T2*-weighted 2D echo-planar images were acquired using a third-party multiband sequence^47,48^, with the following parameters: 235 time points, 36 slices, 2.3 mm isotropic resolution, TR/TE=1000/32.40 ms, flip angle 60°, base resolution: 84 × 84, 192 mm field of view (FoV), multiband factor 4. A whole-brain EPI volume was also acquired and used in an intermediate step in the spatial registration of the partial functional image with the anatomical image. Both the partial and whole-brain volumes consisted of axial slices which were tilted so that they were parallel to the calcarine sulcus for each participant. The central slice overlapped with the calcarine sulcus. A B0 field-map was also acquired to allow for the correction of geometric distortions due to B0 field inhomogeneity in the EPI data^45^. The total acquisition time for all MRI sequences was about 70 minutes.

### MRI data analysis

#### Pre-processing

Cortical reconstruction was performed on the T1-weighted (T1w) image using FreeSurfer’s recon-all function (FreeSurfer software package^49^ version 6.0, http://surfer.nmr.mgh.harvard.edu/).

FMRI data pre-processing was done using the statistical parametric mapping (SPM) software package (SPM12, Wellcome Trust Centre for Neuroimaging, London, UK, http://www.fil.ion.ucl.ac.uk) in MATLAB. The data consisted of six experimental sessions of pRF mapping. The functional images were spatially realigned to the mean of the whole time-series using rigid-body transformations to correct for head motion. The B0 field map image was used to correct EPI geometric distortions. We then performed slice timing correction and intensity bias correction. The mean fMRI volume was co-registered first to the whole-brain fMRI in an intermediary step and then to the anatomical (T1w) image using mutual information.

#### PRF mapping analysis

Population receptive field mapping was done using the SamSrf 9 toolbox for population receptive field (pRF) modelling^50^. The procedure includes projecting the pre-processed retinotopic mapping fMRI data onto the cortical surface and then fitting a standard Gaussian 2d pRF model to the data. The cortical surfaces were the outputs of FreeSurfer’s *recon-all*.

PRF mapping was done by concatenating the imaging data from the six sessions. An occipital cortex ROI was used to restrict the number of surface vertices considered by the mapping algorithm. We used the outputs of the pRF mapping procedure – maps of eccentricity, polar angle, tuning width (standard deviation of the Gaussian) as a measure of pRF size, and variance explained (R^2^) – for further analysis.

We delineated visual area ROIs (V1 dorsal and ventral – V1d and V1v, respectively, V2d/, V3d/v, V4) based on reversals in the polar angle map and restricted our ROIs to realistic eccentricities (i.e., those which were mapped with the stimulus: 0 to 8°) using the eccentricity maps.

### Data analysis

#### Illusion magnitudes

Analyses of illusion data were performed in R (R Development Core Team, 2018). As a measure of intra-rater reliability, we computed two-way mixed effects models (intraclass correlations of type (3,1) or ICC_3,1_) between the two adjustments of each condition^51,52^. More than a simple correlation, ICCs also reflect the agreement between measurements. As suggested by Gignac and Szodorai, we considered correlation coefficients of 0.1, 0.2, and 0.3 as small, medium, and large, respectively^53^. Note, however, that Cohen suggested to consider 0.1, 0.3, and 0.5 as the cut-off values for small, medium, and large effect sizes^54^.

The illusion magnitude (averaged across two trials) was expressed as a difference compared to the reference, except for the HC and EX conditions, where the extracted value was the area of the adjusted ellipse. For each participant and each condition, we combined both conditions of each illusion, i.e., we added the absolute value of the condition, which is usually under-adjusted, to the other condition. Correlations were then computed between each pair of illusions to inspect individual differences in the perception of visual illusions.

#### Visual surface area

In a first analysis, we calculated the surface area of visual areas V1, V2, V3, and V4 within eccentricities bounded by the foveal representation and 8 deg. We then estimated correlations between illusion magnitude and ROI surface area for each illusion-ROI pair. We detected outliers based on the absolute modified *z*-score, which is more robust than the common *z*-score because it is based on the median and median absolute deviation instead of the mean and standard deviation. Any data point which had an absolute modified *z*-score above 3.5 in terms of either illusion magnitude or ROI surface area was removed, as suggested by Iglewicz and Hoaglin^55^.

We used the open-source JASP software^56^ (Jeffreys’s Amazing Statistics Program; https://jasp-stats.org) for Bayesian and frequentist correlation analysis of illusion magnitudes and ROI surface areas. We considered correlation coefficients of 0.1, 0.3, and 0.5 as small, medium, and large effect sizes, respectively^54^. We considered a BF_10_ between 3 and 10 to correspond to substantial evidence in favor of the alternative hypothesis^17^ (in this case: substantial correlation between the two variables).

#### PRF size

The previously mentioned studies that found correlations between size illusion magnitude and V1 surface area argued that their findings could be explained by the relation between subjective size perception and the spatial spread of neural activity^3–5^. While our above analysis used functionally defined surface area as a measure of central cortical magnification across the visual field, we wanted to investigate the role of local spatial spread of neural activity.

Thus, in our second analysis, we extracted pRF sizes at illusion-relevant eccentricities and computed correlations between these pRF sizes and the corresponding illusion magnitudes. This analysis can only be done for illusions in which the targets are at a fixed distance from one another. Moreover, the distance between the targets can only be reasonably determined for circular targets. Therefore, here we focused on the Delboeuf and Ebbinghaus 2 illusions. In both cases, we assumed that participants used one of two strategies: (1) focusing their gaze at the midpoint location between the two targets, or (2) focusing their gaze directly on one or the other of the targets.

In the DB illusion, the distance between the centers of the two targets was 3.92°, with the targets positioned diagonally from each other – one in the upper left and the other in the lower right. Thus, we extracted pRF sizes from the following eccentricities and ROIs: right ventral (corresponding to the top left target) and left dorsal (corresponding to the bottom right target) ROIs, at eccentricities of 3.92° and 1.96° (=3.92°/2, i.e., the midpoint between the two target locations). Similarly, for the EB2 illusion, we considered right and left hemisphere ROIs (since targets were positioned horizontally from each other), at eccentricities of 9.3° (distance between the targets) and 4.65° (midpoint between the targets).

We calculated Bayesian and frequentist correlations between DB and EB2 illusion magnitudes and pRF sizes at the aforementioned eccentricities for each ROI. We removed any data point which had a modified *z*-score above 3.5 in terms of either illusion magnitude or pRF size^55^. As above, we considered correlation coefficients of 0.1, 0.3, and 0.5 as small, medium, and large effect sizes, respectively, and a BF_10_ between 3 and 10 as substantial evidence in favor of the alternative hypothesis.

#### Slope and intercept of pRF size as a function of eccentricity

The above pRF analysis was limited to just two illusions due to the difficulty of identifying “relevant” eccentricities for most illusions in a free-viewing condition and with non-circular targets. To circumvent this issue and characterize the relationship between pRF size and illusion magnitude for all tested illusions, we conducted a final analysis, in which we estimated the slope and intercept of pRF size as a function of eccentricity. An association between illusion magnitude and the slope would indicate that interindividual susceptibility to the given illusion is predicated on the rate at which pRF size increases with eccentricity. Meanwhile, an association between illusion magnitude and the intercept would imply that the parafoveal pRF size is the determining factor for illusion susceptibility.

For each participant, each ROI and the two hemispheres separately, we fit a linear regression model to pRF size as a function of eccentricity, weighted by the pRF model’s goodness-of-fit (R^2^) at each given eccentricity. Fitting was done for eccentricities between 1 and 8° to remove edge-of-stimulus effects (see **Figure S3** for an example). We computed the slope and intercept of the least squares line. We then averaged the slopes and intercepts from the two hemispheres. For some participants, the slope was estimated to be negative due to noise in the data (see **Figure S4**). If this was the case in one hemisphere, we only considered the slope from the other hemisphere. If it was the case in both hemispheres, we omitted the data point from further analysis.

For each illusion and each ROI, we conducted Bayesian linear regression analyses, with illusion magnitudes as the dependent variable and slope and/or intercept as the covariate(s). We tested four models: (1) model that included the slope, (2) model that included the intercept, (3) model that included both slope and intercept, (4) null model that did not include either slop or intercept. We estimated the Bayes Factors (BF_10_) in favor of each of the three alternative models over the null model. As above, we considered a BF_10_ between 3 and 10 to correspond to substantial evidence in favor of the alternative model.

