## Supplementary Information for "Is there a neural common factor for visual illusions?"

**Table S1.** Intra-rater reliabilities expressed as intraclass coefficients (ICC) for both reference-dependent conditions of each illusion (N = 30). *P*-values were not corrected for multiple comparison (i.e., $\alpha = 0.05$).

| Illusion | Condition | ICC coef. | 95% CI | *df* | *p* |
|---|---|---|---|---|---|
| BS | horizontal | 0.622 | [0.394, 0.777] | [29, 29] | <0.001 |
| | vertical | 0.540 | [0.285, 0.723] | [29, 29] | 0.001 |
| CS | left | 0.274 | [-0.029, 0.531] | [29, 29] | 0.068 |
| | right | 0.149 | [-0.159, 0.430] | [29, 29] | 0.212 |
| DB | left | 0.478 | [0.207, 0.681] | [29, 29] | 0.003 |
| | right | 0.597 | [0.360, 0.761] | [29, 29] | <0.001 |
| EB1 | small | 0.784 | [0.632, 0.878] | [29, 29] | <0.001 |
| | large | 0.304 | [0.004, 0.554] | [29, 29] | 0.048 |
| EB2 | small | 0.608 | [0.376, 0.769] | [29, 29] | <0.001 |
| | large | 0.686 | [0.486, 0.818] | [29, 29] | <0.001 |
| EX | black | 0.763 | [0.599, 0.865] | [29, 29] | <0.001 |
| | white | 0.906 | [0.832, 0.948] | [29, 29] | <0.001 |
| HC | black | 0.921 | [0.858, 0.957] | [29, 29] | <0.001 |
| | white | 0.840 | [0.721, 0.911] | [29, 29] | <0.001 |
| ML | outward | 0.528 | [0.270, 0.715] | [29, 29] | 0.001 |
| | inward | 0.658 | [0.445, 0.800] | [29, 29] | <0.001 |
| PD | left | 0.394 | [0.105, 0.621] | [29, 29] | 0.014 |
| | right | 0.699 | [0.505, 0.826] | [29, 29] | <0.001 |
| PZ | down | 0.713 | [0.525, 0.835] | [29, 29] | <0.001 |
| | up | 0.654 | [0.440, 0.798] | [29, 29] | <0.001 |

| | | | | | |
|---|---|---|---|---|---|
| PZh | down | 0.850 | [0.737, 0.916] | [29, 29] | <0.001 |
| | up | 0.642 | [0.422, 0.790] | [29, 29] | <0.001 |
| TT | left | 0.340 | [0.044, 0.581] | [29, 29] | 0.031 |
| | right | 0.511 | [0.249, 0.704] | [29, 29] | 0.002 |
| ZN | left | 0.534 | [0.278, 0.720] | [29, 29] | 0.001 |
| | right | 0.773 | [0.615, 0.871] | [29, 29] | <0.001 |

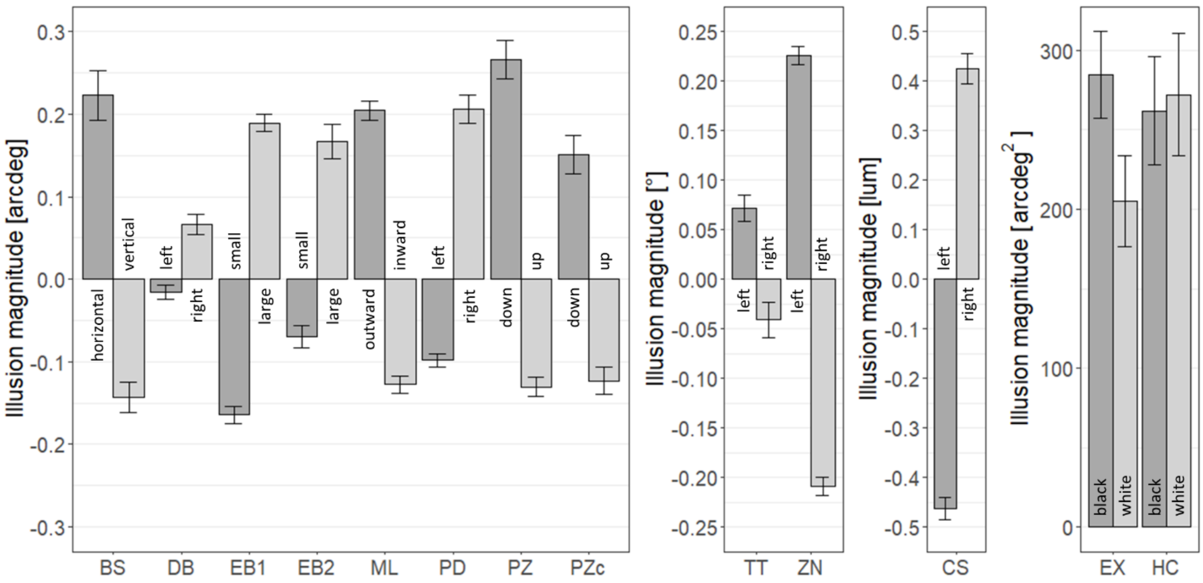


**Figure S1.** Illusion magnitudes for both reference-dependent conditions of each illusion. For all but the EX and HC illusions, positive and negative magnitudes indicate over- and under-adjustments compared to the reference, respectively. Error bars correspond to the standard error of the mean (SEM).

**Table S2.** Illusion magnitudes are expressed as a difference compared to the reference, except in the EX and HC conditions, where we extracted the area in which all shapes were perceptible to the participants. Illusion magnitudes are summarized as mean (*M*), standard deviation (*SD*), standard error of the mean (*SEM*), and 95% confidence interval (95% *CI*) for both reference-dependent conditions of each illusion (N = 30).

| Illusion | Condition | *M* | *SD* | *SEM* | 95% *CI* | Unit |
|---|---|---|---|---|---|---|
| BS | horizontal | 0.223 | 0.163 | 0.030 | 0.061 | arcdeg |
| | vertical | -0.143 | 0.101 | 0.018 | 0.038 | |
| CS | left | -0.462 | 0.123 | 0.022 | 0.046 | luminance |
| | right | 0.425 | 0.166 | 0.030 | 0.062 | |
| DB | left | -0.015 | 0.047 | 0.009 | 0.017 | arcdeg |
| | right | 0.066 | 0.067 | 0.012 | 0.025 | |
| EB1 | small | -0.164 | 0.058 | 0.011 | 0.022 | arcdeg |
| | large | 0.189 | 0.056 | 0.010 | 0.021 | |
| EB2 | small | -0.070 | 0.074 | 0.013 | 0.027 | arcdeg |
| | large | 0.167 | 0.114 | 0.021 | 0.043 | |
| EX | black | 284.659 | 148.808 | 27.169 | 55.566 | $arcdeg^2$ |
| | white | 204.984 | 158.165 | 28.877 | 59.060 | |

| | | | | | | |
|---|---|---|---|---|---|---|
| HC | black | 261.930 | 186.635 | 34.075 | 69.691 | arcdeg$^2$ |
| | white | 272.082 | 210.487 | 38.429 | 78.597 | |
| ML | outward | 0.205 | 0.064 | 0.012 | 0.024 | arcdeg |
| | inward | -0.128 | 0.057 | 0.010 | 0.021 | |
| PD | left | -0.098 | 0.046 | 0.008 | 0.017 | arcdeg |
| | right | 0.206 | 0.093 | 0.017 | 0.035 | |
| PZ | down | 0.266 | 0.126 | 0.023 | 0.047 | arcdeg |
| | up | -0.130 | 0.064 | 0.012 | 0.024 | |
| PZh | down | 0.151 | 0.127 | 0.023 | 0.048 | arcdeg |
| | up | -0.123 | 0.091 | 0.017 | 0.034 | |
| TT | left | 0.072 | 0.071 | 0.013 | 0.027 | ° |
| | right | -0.041 | 0.098 | 0.018 | 0.037 | |
| ZN | left | 0.226 | 0.050 | 0.009 | 0.019 | ° |
| | right | -0.209 | 0.050 | 0.009 | 0.019 | |

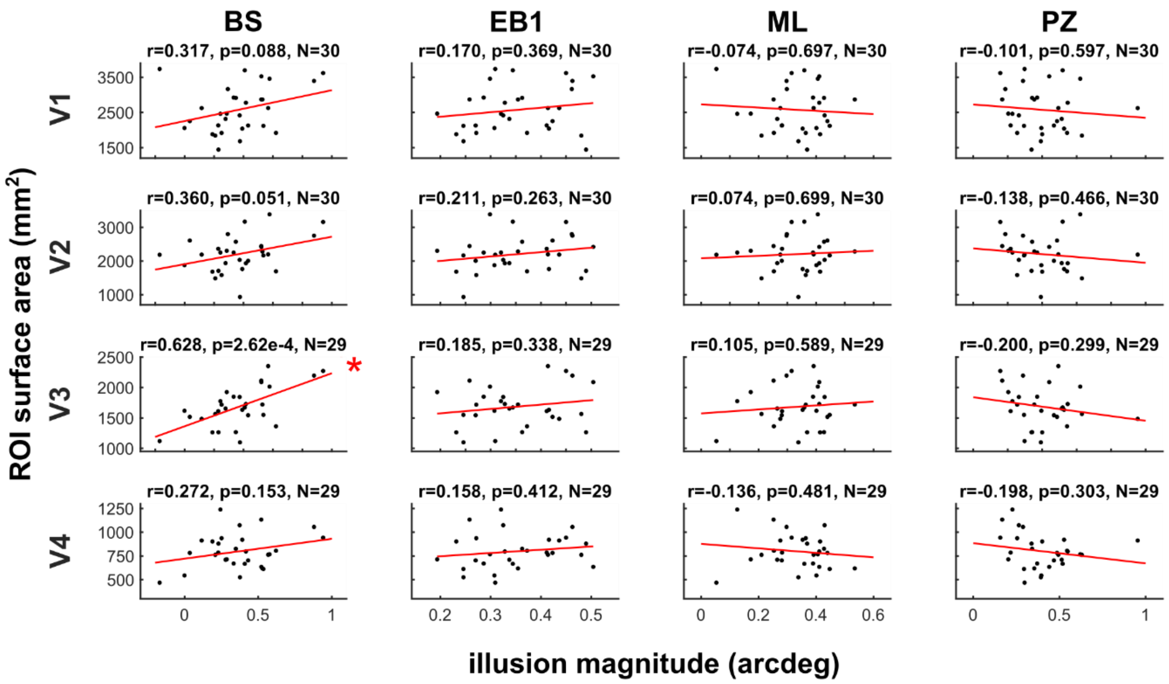


**Figure S1.** Correlations between ROI surface area and visual illusion magnitudes for other size illusions. BS – bisection, EB1 – Ebbinghaus variant 1, ML – Müller-Lyer, PZ – Ponzo. Asterisks mark correlations that are significant ($p < 0.05$, uncorrected). Outliers (modified *z*-score > 3.5) removed. All illusion magnitudes are expressed in units of arcdeg.

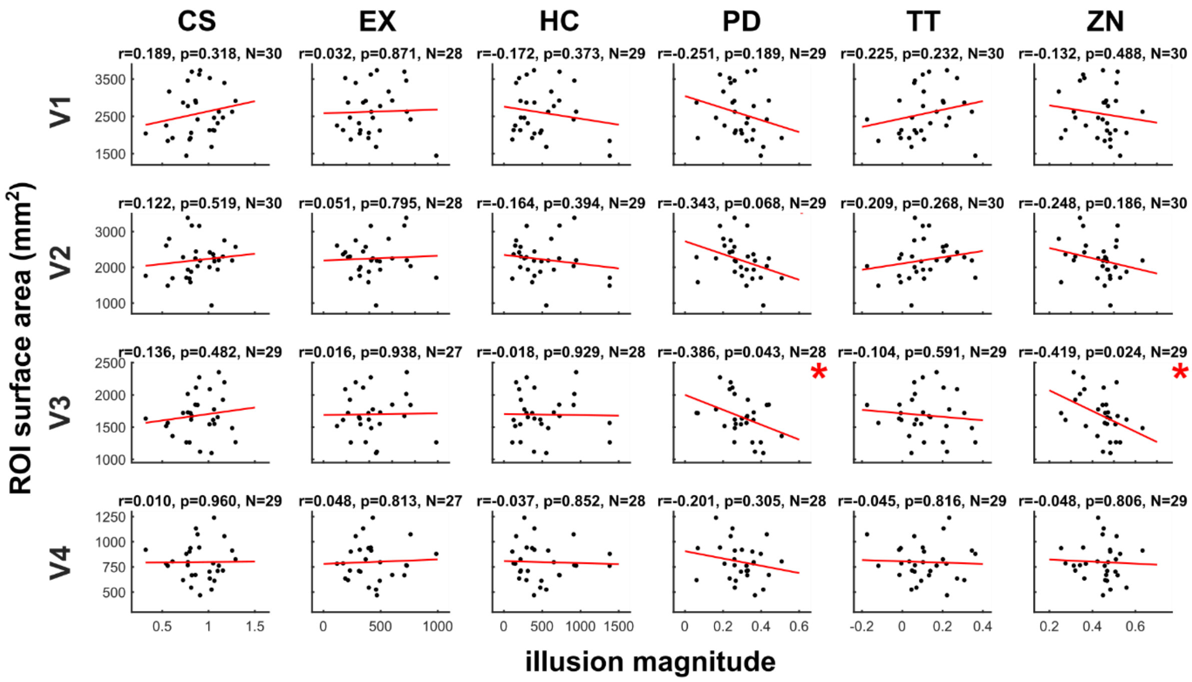


**Figure S2.** Correlations between ROI surface area and visual illusion magnitudes for remaining illusions (uniform textures, contrast, perceived orientation). CS – contrast, EX – Extinction, HC – Honeycomb, PD – Poggendorff, TT – tilt, ZN – Zoellner. Asterisks mark significant correlations ($p$<0.05, uncorrected). Outliers (modified z-score > 3.5) removed. Illusion magnitudes are expressed in different units for different illusions (see Table S2).

**Table S3.** Results of Bayesian linear regression analysis of illusion magnitudes (dependent variable), using slope and intercept of pRF size as a function of eccentricity as covariates. Results consist of Bayes Factors ($BF_{10}$) of three alternative models w.r.t. the null model: (1) model that includes the slope, (2) model that includes the intercept, (3) model that includes both slope and intercept. Instances of substantial evidence in favor of either the null or the alternative model (BF ∈ [3,10) are marked in bold with an asterisk. Instances of substantial evidence in favor of the null hypothesis ($BF_{10}$ ∈ [0.1,0.333), corresponding to $BF_{01}$ ∈ [3,10)) are marked in red and bold with a dagger (†). Outliers (modified z-score > 3.5) removed.

| Visual area | Illusion | N | $BF_{10}$ | | |
|---|---|---|---|---|---|
| | | | slope | intercept | slope+intercept |
| V1 | BS | 30 | 0.384 | 0.404 | 0.334 |
| | CS | 30 | 0.359 | 0.528 | **0.260†** |
| | DB | 30 | 0.545 | 1.377 | 0.580 |
| | EB1 | 30 | 0.354 | 0.346 | **0.168†** |
| | EB2 | 29 | 0.515 | 0.380 | **0.246†** |
| | EX | 28 | 0.916 | 0.792 | **0.103†** |
| | HC | 29 | 0.765 | 0.373 | 0.386 |
| | ML | 30 | 0.442 | 0.406 | **0.208†** |
| | PD | 29 | 0.464 | 0.481 | 0.991 |
| | PZ | 30 | 0.349 | 0.516 | **0.270†** |
| | PZh | 28 | 1.161 | 0.653 | 0.507 |
| | TT | 30 | 0.398 | 0.517 | 0.706 |
| | ZN | 30 | 0.487 | 0.345 | **0.301†** |
| V2 | BS | 30 | 0.901 | 0.368 | 0.517 |
| | CS | 30 | 0.526 | 0.352 | **0.276†** |
| | DB | 30 | 0.431 | 0.880 | 0.403 |
| | EB1 | 30 | 0.356 | 0.344 | **0.174†** |

| | | | | | |
|---|---|---|---|---|---|
| | EB2 | 29 | 0.350 | 0.402 | **0.234†** |
| | EX | 28 | 0.476 | 0.356 | 0.337 |
| | HC | 29 | 0.640 | 0.356 | 0.639 |
| | ML | 30 | 0.360 | 0.351 | **0.171†** |
| | PD | 29 | 0.370 | 1.082 | 2.523 |
| | PZ | 30 | 0.366 | 0.350 | **0.201†** |
| | PZh | 28 | 0.428 | 0.379 | **0.205†** |
| | TT | 30 | 0.663 | 0.363 | 0.916 |
| | ZN | 30 | 0.350 | 0.415 | **0.262†** |
| V3 | BS | 30 | 0.677 | 1.565 | 0.669 |
| | CS | 30 | 0.358 | 0.395 | **0.332†** |
| | DB | 30 | 0.572 | 1.984 | 0.998 |
| | EB1 | 30 | 0.351 | 0.345 | **0.169†** |
| | EB2 | 29 | 0.350 | 0.488 | 0.430 |
| | EX | 28 | 0.353 | 0.354 | **0.174†** |
| | HC | 29 | 0.355 | 0.349 | **0.182†** |
| | ML | 30 | 0.347 | 0.585 | 0.469 |
| | PD | 29 | 2.331 | 1.853 | 1.074 |
| | PZ | 30 | 0.358 | 0.371 | **0.176†** |
| | PZh | 28 | 0.378 | 0.361 | **0.184†** |
| | TT | 30 | 0.359 | 0.364 | **0.173†** |
| | ZN | 30 | 0.408 | 0.411 | **0.196†** |
| V4 | BS | 30 | 0.471 | 0.344 | **0.307†** |
| | CS | 30 | 0.356 | 0.525 | **0.274†** |
| | DB | 30 | 0.412 | 0.368 | **0.194†** |
| | EB1 | 30 | 0.387 | 0.393 | 0.365 |
| | EB2 | 29 | 0.782 | 0.348 | 0.809 |
| | EX | 28 | 0.359 | 0.438 | **0.225†** |
| | HC | 29 | 0.352 | 0.388 | **0.228†** |
| | ML | 30 | 0.940 | 0.650 | 0.416 |
| | PD | 29 | 0.348 | 0.349 | **0.168†** |
| | PZ | 30 | 0.344 | 0.415 | **0.228†** |
| | PZh | 28 | 0.689 | 0.370 | 0.395 |
| | TT | 30 | 0.344 | 0.350 | **0.168†** |
| | ZN | 30 | 1.016 | 0.918 | 0.501 |

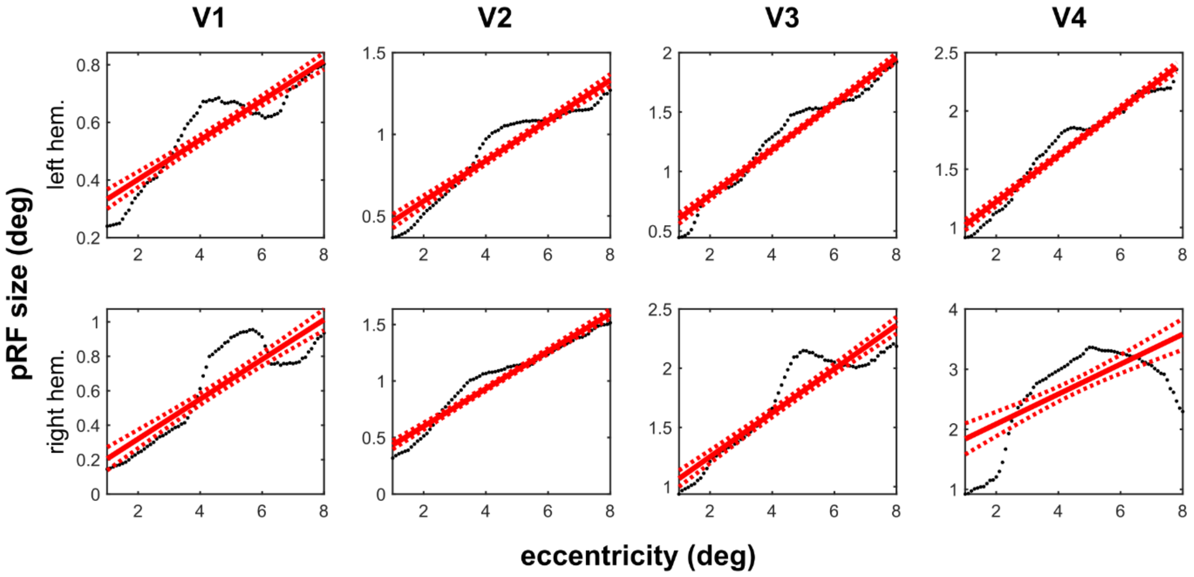


**Figure S3.** Linear regression model fit to pRF size as a function of eccentricity, weighted by the pRF model's goodness-of-fit ($R^2$), across ROIs and in the two hemispheres for a sample participant. All slopes are positive, as would be expected based on known properties of retinotopic organization. Dotted red lines represent the 95% confidence bounds.

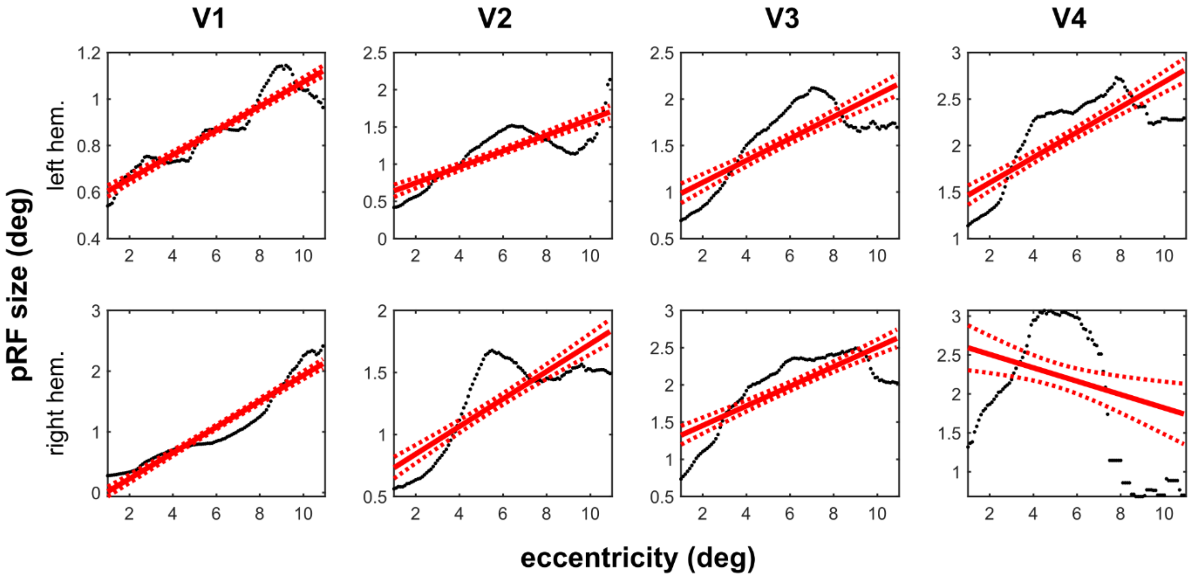


**Figure S4.** Linear regression model fit to pRF size as a function of eccentricity, weighted by the pRF model's goodness-of-fit ($R^2$), across ROIs and in the two hemispheres for another sample participant. Here, due to noisy pRF estimates in higher eccentricities, slope is estimated to be negative in V4 of the right hemisphere. Such data points were discarded. Dotted red lines represent the 95% confidence bounds.

***Population receptive field mapping stimulus***

The wedge systematically stimulated different polar angles segments of the visual field, rotating around fixation in 60 discrete steps of 6°. The wedge completed three cycles per scanning session. Independently, the ring systematically stimulated different eccentricity bands by expanding and contracting consecutively. The ring completed five cycles per session. The steps of the wedge and ring were synchronized with the start of each volume of the fMRI acquisition, i.e., they lasted 1 s each. The wedge and ring aperture revealed a ripple background texture (as previously described[1,2]), which changed every 500 ms. The ripple background consisted of black and white patterns, with a contrast of 100%. Outside of the aperture, the background of the display was set to medium gray.

We included six sessions. In odd scanning sessions, the wedge rotated clockwise, and the ring had an expanding starting direction. In even sessions, this was reversed, with a counterclockwise ring rotation and contracting wedge starting direction.

The wedge stimulus was presented from 0.07° from fixation to an eccentricity of 12°, with an angular size of 12° at all eccentricities. As in previously referenced work[2], the diameter of the outer boundary of the ring was determined as a function of stimulus steps *x*:

$$f(x) = 17e^{\left(-4+\frac{1}{9}x\right)}$$

The stimulus steps $x$ increased or decreased with each acquired volume in steps of 0.9 from 1.5 to 33, yielding 36 discrete steps. The minimum eccentricity of the inner boundary of the ring stimulus was 0.25°; otherwise, the radius of the inner boundary was equal to 75% of the outer boundary.

Participants were instructed to always look at the fixation point (0.13° in diameter). To aid fixation, participants performed a simple fixation task. The fixation point changed color (red, green, white, black in pseudo-random order) with a stimulus onset asynchrony (SOA) of 3±1 s. Participants were asked to push a button within 1.5 s after each color change. The task was designed to keep participant’s gaze at the fixation point. Additionally, low-contrast crosshairs were presented to aid fixation. The crosshairs consisted of 12 radial lines from the fixation dot to the edge of the screen and 8 concentric circles, centered on fixation and increasing in radius in steps of 1.5°, starting from 2°.

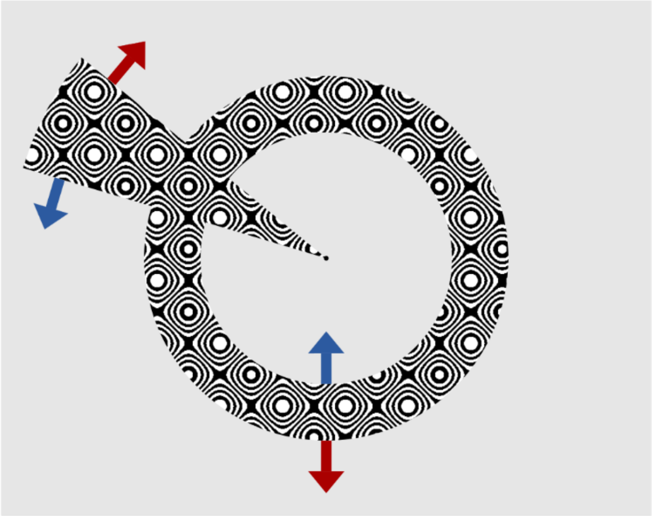

**Figure S5. PRF mapping stimulus.** The stimulus consisted of a rotating wedge and an expanding and contracting ring aperture, which reveals a ripple texture background. Six sessions of pRF mapping were included in the experiment. In each session, the wedge completed three cycles and the ring completed five cycles. During odd-numbered sessions, the wedge rotated clockwise, and the ring started out by expanding (indicated by red arrows). During even-numbered sessions, the wedge rotated counterclockwise, and the ring started out by contracting (indicated by blue arrows).
